# Chip-Based 3D Interferometric Nanoscopy

**DOI:** 10.1101/2025.05.13.653677

**Authors:** Wei Wang, Hangfeng Li, Yilin Wang, Samuel F. H. Barnett, Zengxin Huang, Pakorn Kanchanawong

## Abstract

Ultra-high resolution 3D single-molecule localization microscopy (SMLM) traditionally requires complex dual-objective lenses (4Pi) configurations to enhance axial (z) precision through interferometry. Here we present a streamlined chip-based alternative, Silicon-assisted interferometric Localization Microscopy (SiLM), which achieves comparable performance using a single-objective lens design. By combining tunable axial structured illumination field, arising from surface-generated excitation interference, with asynchronous interferometry, SiLM enhances axial localization precision to approximately twice that of the lateral (xy), comparable to 4Pi-based methods. Additionally, SiLM provides intrinsic axial self-referencing, offering dramatically improved robustness against mechanical drift. Our method is readily implementable on standard SMLM-capable microscopes and supports a broad range of applications including dual-color imaging, extended-depth imaging, and live-cell 3D single-molecule tracking. Using SiLM, we demonstrate accurate mapping of the stratified nanoscale architecture of integrin-based focal adhesions, establishing it as a powerful and accessible method for high-precision 3D structural cell biology.

## Introduction

Diffraction limits the spatial resolution of light microscopy to typically around 200-300 nm in the lateral (x, y) dimension and multiple-fold poorer along the axial (z) dimension^1^. Widely used methods to surpass these limitations such as Single Molecule Localization Microscopy (SMLM) depend on the precise localization of individual fluorescent molecules^2-5^. However, while SMLM can achieve molecular-scale precision in the lateral dimension^6^, attaining comparable axial precision remains an ongoing challenge^7^. Among 3D SMLM strategies, the modification of the point spread function (PSF) such as by astigmatism or PSF engineering^8,9^ offers a modular approach, but still suffers from axial precision being 2-3 times worse than the lateral. This disparity limits their effectiveness in applications that demand precise axial measurements such as mapping molecular-scale organization within complex cellular structures. Interferometric SMLM (iSMLM) techniques which use dual-opposed objective lenses in 4Pi configuration, such as iPALM, 4Pi-SMS, and 4Pi-STORM exploit fluorescence self-interference to significantly improve axial precision to 2-3 times better than the lateral precision^10-12^. This enabled iSMLM to theoretically approach the fundamental quantum-limited precision^13,14^. However, while iSMLM techniques have been instrumental in elucidating major biological insights^15,16^, the demanding optical instrumentation and laborious operation limit their accessibility and experimental throughput. To address this, we therefore sought to develop a 3D nanoscopy technique capable of comparable performance but with simplified and flexible operation.

In recent years, super-resolution microscopy methods that employed various forms of structured excitation in conjunction with single-molecule analysis have achieved dramatic gain in imaging precision. Beyond MINFLUX^17^, these include several modulated excitation SMLM approaches. For example, in ROSE^18^ and SIMFLUX^19^, lateral interference patterns were utilized to achieve a two-fold lateral precision improvement. Axial precision enhancement was demonstrated in ROSE-Z, which used an excitation standing wave generated by two counter-propagating illumination^20^. Conversely, by using time-modulated tilted excitation, ModLoc demonstrated ultra-deep axial range, albeit with similar axial precision compared to the lateral^21^. By relying on the interference of coherent laser illumination, the fluorophore z-position relative to the axial interference fringe can be decoded from the temporal intensity sequences. Due to their dependence on temporal excitation modulation and the millisecond blinking kinetics of SMLM fluorophores, sophisticated fast-switching optical components are required. Nevertheless, these advances suggest that the modulated excitation approach could provide a basis for a simplified single-objective lens design with axial precision comparable to iSMLM.

Structural analysis of multi-component biomolecular complexes requires precise alignment of a large number of datasets to a common reference frame. This requires precise registration using various forms of internal reference such as the substrate plane, structural landmarks, or internal object symmetry where applicable^22^. The aggregate precision therefore depends on both the precision of the 3D nanoscopic imaging and the precision of dataset registration. For axial super-resolution analysis, the z-coordinate is typically extracted in relation to the calibration model. Techniques that rely on surface-generated processes such as evanescent field^23-25^ or metal-induced energy transfer^26-28^, provide axial coordinate referenced to the surface which facilitates both accurate and precise axial coordinate determination. In contrast, axial interference fringes used in previous iSMLM or modulated excitation SMLM methods are mechanically uncoupled from the specimen, hence entailing sensitivity to drift and the need for additional alignment between datasets. These considerations suggest that a 3D nanoscopy method based on self-referenced axial interference fringes should provide high accuracy, high precision, and high stability measurements, beneficial for streamlining or upscaling structural cell biology investigations.

In this study, we introduce Silicon-assisted Interferometric Localization Microscopy (SiLM) which provides ultra-high 3D imaging precision using a simple single-objective lens design. By using Silicon chip as a sample-integrated phase-shifter, this enables the generation of tunable axial interference fringes analogous to iSMLM, despite utilizing only a single objective lens. Interferometric 3D localization of blinking fluorophores is demonstrated using synchronized modulation of excitation incidence angle and spatial multiplexing, achieving ∼2X better axial precision compared to the lateral, comparable to iSMLM performance. Furthermore, due to self-referenced axial interferometry, the axial interference fringes generated in SiLM are highly resistant to mechanical drift. This enables robust and stable z-coordinate determination. The capabilities of SiLM are demonstrated by resolving the architecture of microtubule filaments and actin cortex, as well as multi-color imaging, extended-depth imaging, and live-cell 3D single-molecule tracking. Structural cell biology application of SiLM is highlighted by mapping the nanoscale architecture of integrin-based focal adhesions, showing a remarkable correspondence with prior iSMLM results^29^. SiLM thus represents a high-performance, cost-effective, and versatile approach that can be readily applied to most existing SMLM-capable microscopy platforms to enable access to advanced 3D nanoscopy capability.

## Results

### Integrating Surface-generated Axial Structured Illumination with Single-Molecule Localization Microscopy

Recent chip-based nanoscopy methods utilize mass-produced chip as imaging substrates to simplify the optical complexity of the imaging systems, but are still limited to 2D super-resolution capability^30,31^. Instead of waveguide, specular reflection is effective in generating axial interference, essential for enhancing z-axis precision^32-36^. We are particularly inspired by previous axial structured illumination microscopy techniques, such as VIAFLIC^37,38^ and SAIM^39^, which offer high precision in axial (z) dimension comparable to iSMLM methods^16^. Furthermore, the z-coordinates obtained by these techniques are stringently referenced against the Silicon chip substrate of each specimen, greatly facilitating analysis of biological structures since additional axial registration is not required. However, the lack of lateral (xy) super-resolution capability limited their application potential to specialized use cases^15,16,40,41^. Here we show that by integrating these approaches with asynchronous interferometry, accurate and precise 3D nanoscopy, comparable to iSMLM benchmark, can be achieved with a much streamlined optics.

As illustrated in **Fig. 1a-c**, the excitation standing wave generated above the reflective surface depends on both the incidence angle and the excitation wavelength, which can be analytically calculated using Fresnel equations (see Methods, **Fig. S1**). The period and phase of the standing wave varies with the excitation wavelength, incidence angles, refractive indices, and thickness of the optical layers (**Fig. S1d-g**). For interferometric axial coordinate determination, the phase of the excitation standing wave should be shiftable over the full 2π range. As shown in **Fig. S1f-g**, this requires axial separation of the specimen of at least ∼λ/2n above the specular surface. In this study, we use Silicon chips with ∼500 nm thermal SiO_2_ layer which provides a biocompatible surface for direct cell culture^39^. The optical configuration outlined in **Fig. 1a** and **Fig. S1a-b** can be easily implemented on an inverted microscope, whereby the Si chip is mounted in aqueous media, facing downward in a glass-bottom dish. As such, a fluorophore located at a specific z-position above the SiO_2_ surface encounters different phases of the standing wave as the incidence angles are varied, leading to a variation in fluorescence emission intensity (**Fig. 1b**) that encodes z-coordinate information. Since the SiO_2_ layer acts as a built-in phase shifter, its thickness is a critical parameter that must be determined with nanometer precision^38,39,42^. This can be performed using techniques such as variable incidence angle ellipsometry, available in most material characterization laboratories. The optimal set of incidence angles for achieving high axial precision can be determined numerically as a function of SiO_2_ layer thickness and excitation wavelength (**Supplementary Software**), with parameters for common laser wavelengths tabulated in **Table S1** (see **Supplementary Notes** I). For example, with SiO_2_ thickness of ∼480 nm and excitation wavelength of 639 nm, the incidence angles of 16°, 36°, and 48° are calculated to provide optimal z-precision. **Fig. 1d-e** shows the interference pattern along the axial direction, with a peak-to-valley ratio greater than 10:1 and a modulation period of approximately half the excitation wavelength.

**Figure 1.**
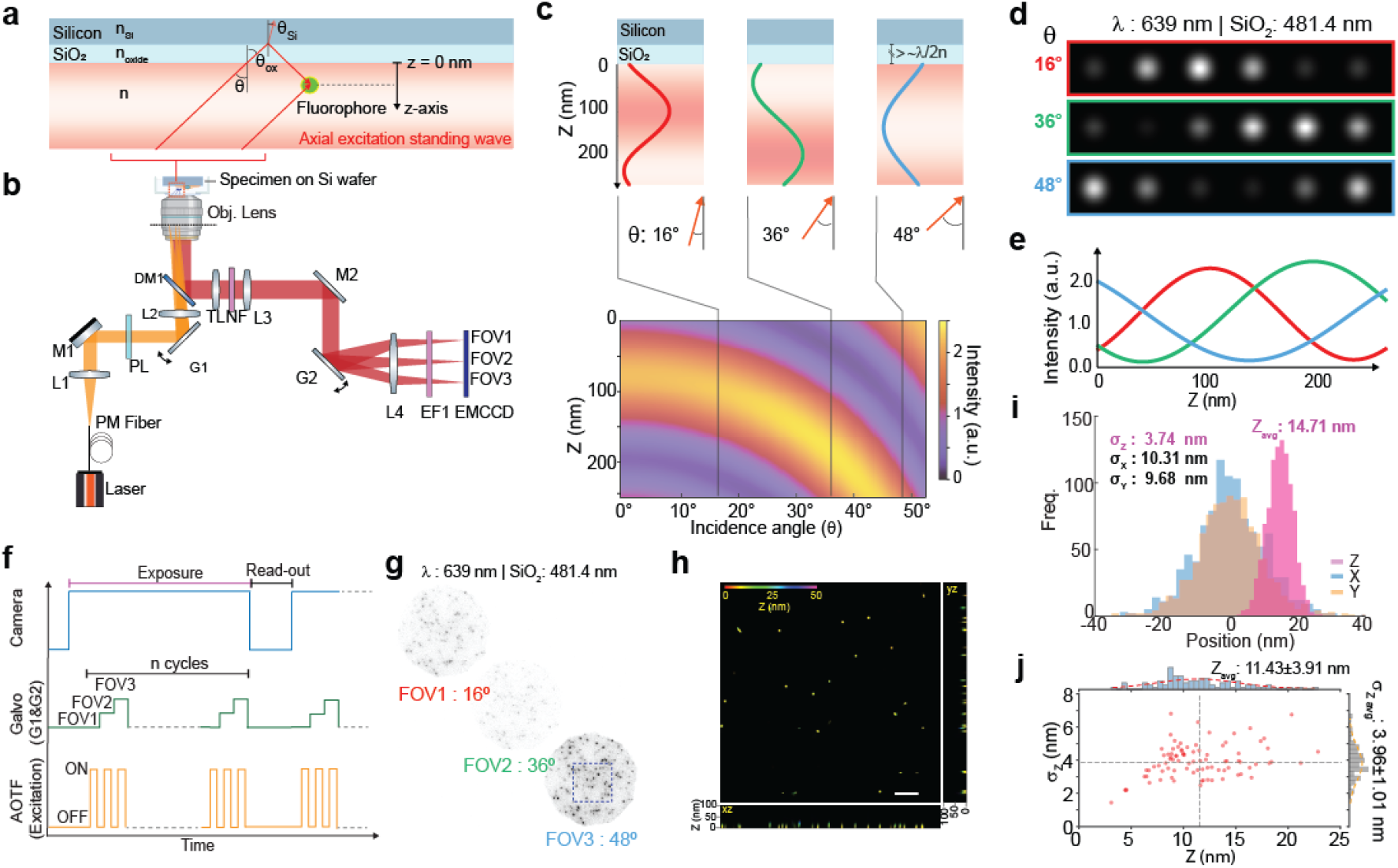
Principles of chip-based interferometric 3D nanoscopy by SiLM. (**a**) Definition of coordinate system (see also Fig. S1b). Fluorophore z-positions are defined relative to the SiO_2_surface of the Si chip (z = 0 nm). The variable incidence angle in aqueous media is defined by θ. (**b**) Schematic diagram for SiLM instrumentation. The specimen on Si chip is placed upside-down in aqueous media in a glass-bottom dish. Galvomirror G1 cycles through different values of θ, while galvomirror G2 synchronously redirects emission to corresponding field-of-view (FOV) on EMCCD camera. See also Fig. S3. (**c-d**) Angle-dependent axial interference fringes for TE-polarized excitation. For 639-nm excitation and 481.4 nm SiO_2_ thickness, the optimal incidence angles are 16*°*, 36*°*, and 48*°*, resulting in standing waves with approximately 120*°* mutual phase differences. (**e**) Simulated fluorophore intensity at θ=16*°*, 36*°*, and 48*°* as a function of z-coordinate. (**f**) Timing diagram depicting the synchronized control of acquisition, camera exposure, galvomirrors, and laser excitation. (**g**) Representative EMCCD frame with 3 FOVs corresponding to θ = 16*°*, 36*°*, and 48*°*. Specimens are fluorescent beads with nominal 27-nm diameter deposited on Si wafer with 481.4 nm SiO_2_ thickness. Intensity variation between FOV corresponding to prediction in (d-e) are observed. (**h**) Reconstructed 3D images corresponding to boxed region in (g), with color encoding z-dimension. Transverse (xz, and yz) projection are depicted with expanded axial scale as indicated in the figure. Scale bar: 1 µm. (**i**) 3D localization precision of single bead from (**g-h**), with xy-coordinates defined relative to bead centroid and z-coordinate relative to SiO_2_ surface. Fits to histograms and σ of the fits are indicated, showing ∼2X better axial precision relative to the lateral. (**j**) Composite plot depicting the axial (z)-position and σ_z_ for individual beads from (g-h), and histograms with gaussian fit. Average bead z-position of 11.43 nm±3.91 nm was found, in good agreement with known bead dimension.

The optical schematics of our system, here called SiLM (***S***ilicon-assisted ***i***nterferometric ***L***ocalization ***M***icroscopy), is shown in **Fig. 1b** whereby the excitation path of an inverted microscope is modified by the addition of a scanning galvo mirror (G1) conjugated to the specimen plane to enable rapid shifting of the incidence angle. In the detection path, another scanning galvo mirror (G2) is used to synchronously map output signals to different field-of-views (FOVs) on the EMCCD camera. The timing diagram for the acquisition is shown in **Fig. 1f**, whereby fluorescence signals from multiple cycles of angle-scanning are integrated for each EMCCD read-out frame to minimize effects of asynchronicity (see below). Single-molecule images from each FOV effectively correspond to different incidence angles (**Fig. 1g**). By fitting the single-molecule intensity triplets to the theoretical model, the z-position can be extracted, analogous to 3-phase interferometric analysis used in 4Pi-based iPALM techniques^10^. Fit using three incidence angles is sufficient to achieve homogeneous axial localization precision over a range of approximately 250 nm (or ∼λ/2n), provided that the corresponding standing waves are mutually phase-shifted by approximately 2π/3 (∼120°) (**Fig. S2a-h**). Since the period of the standing waves vary with incidence angles (**Fig. S1f**), phase-wrapping effect inherent to iSMLM^43^ can be avoided by making use of 4-(or more) angles to extend the imaging depth (**Fig S2i-l**, see below). Note that the excitation intensity is also attenuated as a function of incidence angle, which can be theoretically or empirically corrected to further enhance z-coordinate accuracy (**Fig. S1c, S3e-h**, Methods). Meanwhile, the lateral (x, y) coordinates can be determined through standard localization analysis of the summed single-molecule images^2,3^, while 3D super-resolution reconstruction can be carried out once sufficient single-molecule coordinates are accumulated. Importantly, unlike 4Pi-based iSMLM methods^44-46^ which require laborious alignment and calibration, in our method the calibration step is drastically simplified, involving primarily imaging the emission FOVs at a fixed incidence angle to assist in multi-FOV registration.

### Analysis and Validation of SiLM Localization Precision

Numerical analysis of the theoretical precision for SiLM is shown in **Fig. S2b-h**. Benefitting from axial interferometry, in the limit of infinite fluorophore ‘on’ time, whereby the intensities are equally sampled between different FOVs, the axial resolution is calculated to be 2-3X better than the lateral resolution. To validate the resolution capability, we first imaged fluorescent beads with nominal 27-nm diameter deposited on the SiO_2_ surface, using the optimal incidence angles of 16°, 36°, and 48° for SiO_2_ thickness of 481.4 nm and excitation wavelength 639 nm. As shown in **Fig. 1g** and illustrated in **Fig. S3**, consistent with theoretical predictions (**Fig. 1c-e**), FOV-3 (incidence angle: 48°) exhibits maximum intensity, while FOV-2 (incidence angle: 36°) exhibits the lowest intensity. From 3D localization analysis (**Fig. 1h-j,** see analysis flowchart in **Fig. S4**), the axial (z) positions of the beads are centred at ∼11.43 nm above the substrate, in good agreement with the expected bead radius. As shown in **Fig. 1i**, the axial precision is ∼3.74 nm with a mean value of 14.71 nm, approximately 2.5 times better than the lateral precision (10.31 nm and 9.68 nm), for a mean photon count of ∼1574, consistent with our theoretical analysis. Altogether, SiLM achieves both accurate and precise 3D nanoscopic measurements comparable to the much more complex iSMLM methods while relying on a significantly simplified optical configuration and operation.

### Localization Precision of Asynchronous Axial Interferometry

Blinking fluorophores in SMLM typically exhibit ‘on’ time in the range of tens of milliseconds, comparable to the camera exposure duration. This is expected to introduce additional uncertainty in the temporal domain for asynchronous detection schemes used in modulated excitation SMLM techniques^47^, whereby both the fluorophore ‘on’ time and the start time of each blinking molecule are random (**Fig. 2a**). Previous methods sought to mitigate the effect of asynchronicity by the use of high-speed demodulation optical components, such as resonant galvanometers, acousto-optic modulators (AOMs), or optical Pockels cells^20,21^. However, these components require complex control and offers relatively small modulation areas while incurring high cost. Comparatively, the scanning galvo mirrors used in our implementation are cost-effective components that provide simple, flexible, and precise angular control but are limited in terms of scan speed to ∼1 kHz. Thus, we hereby developed a theoretical framework for analyzing the influence asynchronous detection on the precision of axial interferometry.

**Figure 2.**
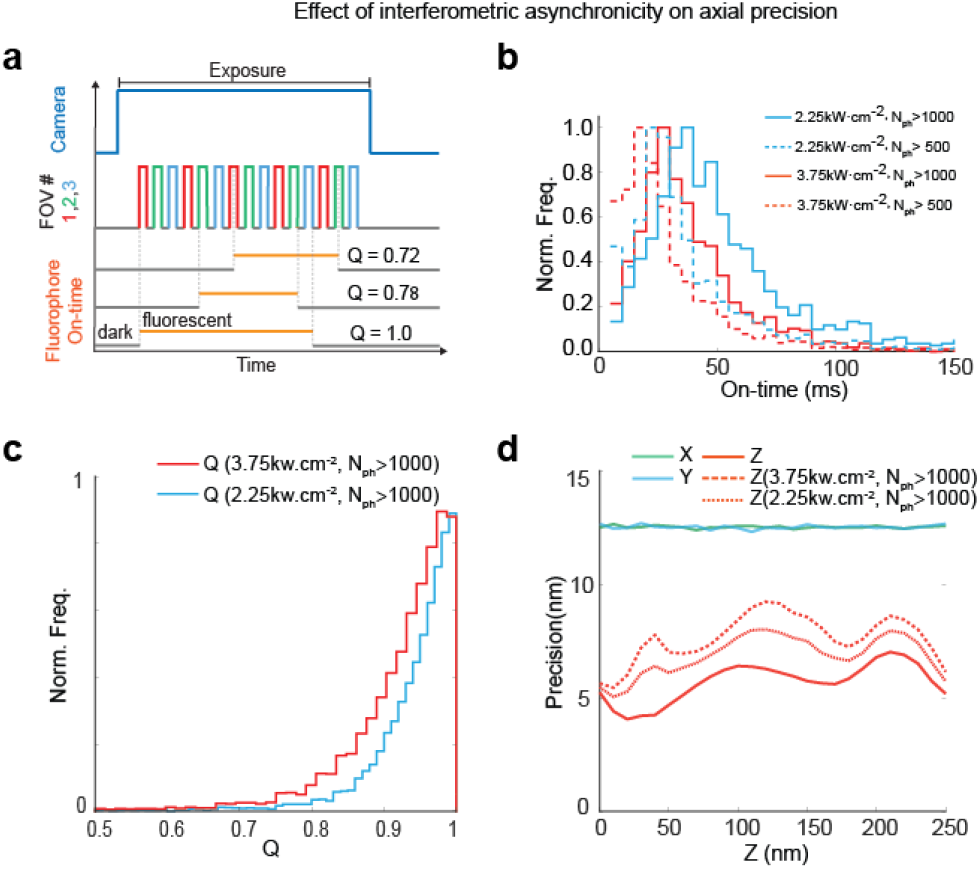
Effect of interferometric asynchronicity on axial precision. (**a**) Stochastic blinking of the fluorophores gives rise to unequal duty ratio among different FOVs in SiLM, described by interferometric quality factor (Q; equal duty ratio among FOVs corresponds to Q = 1). (**b**) Fluorescent ‘on’ time of AlexaFluor 647 under different illumination laser power as a function of photon count (N_ph_) threshold. By optimizing laser intensity, and optimizing N_ph_ threshold, the mean ‘on’ time can be increased. (**c**) Simulated distribution of interferometric asynchronicity under two illumination conditions. (**d**) Theoretical 3D precisions of SiLM, comparing the fully synchronous limit (solid red line) with scenarios where Q are defined as in (c). The axial precision is slightly attenuated but remains much better than the lateral precision (blue, green), which are not affected by Q.

Due to the stochastic and finite fluorophore ‘on’-time, different sampling duty ratio can be expected for each FOV (**Fig. 2a**), which inevitably give rise to the deviation of measured intensity from the theoretical model and degrade axial localization precision. To address this, we defined a parameter Q to represent the interferometric quality factor, defined as mean(t_i_)/max(t_i_) where t_i_ denotes the duration of a blinking molecule in the i^th^ cycle. For the ideal case (Q=1), a constantly ‘on’ fluorophore is equally sampled across all FOVs. To mitigate the impact of asynchronicity, fluorophore ‘on’ time characteristics were first characterized as shown in **Fig. S5e-h**. We then implemented rapid interlacing of FOVs relative to the fluorophore ‘on’ time (**Fig. S3a-d**). We found the acquisition cycle time of 2*n* ms to be optimal for SiLM, where *n* is the number of FOVs, with 1.1 ms allocated for the galvo settling time and 0.9 ms for laser dwell time at each incidence angle/FOV pair (**Fig. 1f, Fig. S3d**). The number of cycles depends on specific fluorophores; for example, 16 cycles per frame is used for Alexa Fluor 647. Additional methods to ensure high Q can be implemented. First, the emission intensity of individual fluorophore is proportional to both the laser intensity and the duration of ‘on’ time. As such, there is a correlation between ‘on’ time and emission photon count, as shown in **Fig. 2b**, implying that high photon count correlates with high Q. Using the excitation pulses defined by **Fig. 1f**, the mean ‘on’ time, τ_on_, of Alexa Fluor 647 was found to be 33 ms when the minimum photon-count cut-off, *N*_*thres*_, is set to 500 photons. For *N*_*thresh*_of 1000 photons, τ_on_ increases to 46.2 ms, reflecting that molecules with shorter ‘on’ times are filtered out. Secondly, this can be complemented by modulating excitation intensity since previous studies have shown that τ_on_ is inversely proportional to the laser intensity^47^. For example, by reducing laser intensity to 60% (2.25kW/cm^2^), τ_on_ for Alexa Fluor 647 becomes 43.8 ms with *N*_*thresh*_of 500 photons, and 63 ms with *N*_*thresh*_of 1000 photons (**Fig. S5e-f**).

To simulate the influence of ‘on’ time on localization precision, each molecule is modelled to be randomly activated and stay emitting for a random duration, *T*_on_. Based on this, the emission duration for each cycle is simulated, and the Q value calculated accordingly (for more details, see **Supplementary Notes II**). As shown in **Fig. 2b**, the distribution of *T*_on_ gives rise to the distribution of Q as shown in **Fig. 2d**. Using a laser intensity of 3.75 kW/cm^2^ and *N*_*thres*_of 1000, Q mostly falls between 0.7 and 1, with further improvement upon reduction in laser intensity. The distribution of 3D localization precision for different Q is shown in **Fig. 2d**. In the ideal case (Q = 1), the axial precision is 2-3 times better than the lateral precision, comparable to 4Pi-based iSMLM methods^10,12,45^. At high laser power, a reduction in Q results in axial precision being 1.4-1.8X better than the lateral precision, reflecting the influence of randomness in the temporal dimension. In contrast, at lower laser power, the axial precision improves to 1.6-2X better than the lateral, averaging to 1.85X across the entire range. As shown in **Fig. S5b-d**, for molecules with an τ_on_ between 42-48 ms, the axial precision is close to the theoretical value. Even with shorter τ_on_ ranging from 12-18 ms, the axial precision is approximately 1.2 to 1.6 times better than the lateral precision. Taken together, our analysis shows that while a small trade-off in axial precision is incurred when using scanning galvo mirror with *ms* switching times, a significant enhancement in axial precision at 1.8X better than the lateral precision is still preserved, indicating that galvo-based asynchronous interferometry could provide a practical and cost-effective solution for 3D nanoscopy. Consistent with this, Fourier Ring Correlation analysis (**Fig. S6a-d**) of experimental datasets yielded a ∼1.8X enhancement of axial resolution relative to the lateral, in good agreement with the theoretical analysis in **Fig. 2d**.

### Imaging 3D cellular ultrastructure by SiLM

To demonstrate the bioimaging capabilities of SiLM, we first imaged various 3D nanoscale structures in cells (imaging parameters in **Table S2**). As SiLM provides the absolute axial positions with respect to SiO_2_ surface, for long-term acquisition, automated focus-correction in commercial microscopy system such as Nikon PerfectFocus System® can maintain sample movement within hundreds of nanometers, which is sufficient to preserve the ultra-high precision of the reconstructed axial position, and only lateral drift needs to be corrected (**Fig. S7**). As shown in **Fig. 3a-h**, the microtubule filaments in fixed and immunostained COS-7 cells are imaged using AlexaFluor 647-conjugated secondary antibody. From isometric cross-section views of parallel microtubules (dashed lines in **Fig. 3a, Fig. 3b**), or contiguous sections of filaments (boxes in **Fig. 3a, Fig. 3c,f**), the ring-like cross-profiles of the microtubules are clearly discerned. The coordinates of the filament cross-sections from **Fig. 3c** and **Fig. 3f** are aligned and plotted in **Fig. 3d** and **3g**, respectively. In **Fig. 3e**, the axial histogram corresponding to **Fig. 3d** exhibits two distinct peaks, separated by 26.09 nm, and with standard deviations of 8.79 and 10.53 nm. Similarly, in **Fig. 3h**, the radial distribution of coordinates from regions in **Fig. 3g** exhibit a peak with standard deviation of 10.26 nm, and the mean radius of the microtubule is centred at 23.32 nm.

**Figure 3.**
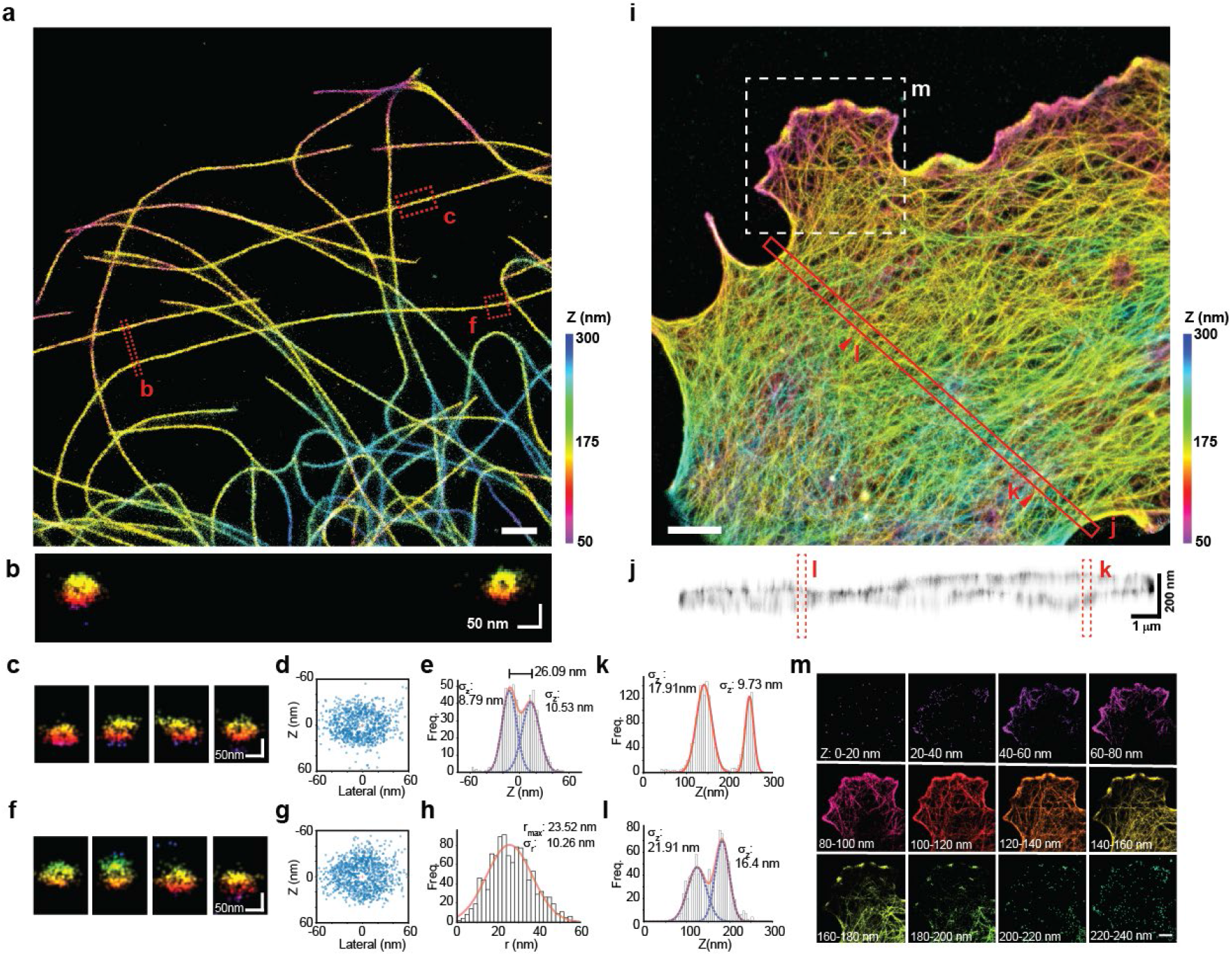
Nanoscopic 3D Bioimaging by SiLM. (**a**) Axial color-coded 3D images of AlexaFluor647-immunolabelled microtubule filaments in COS-7 cells. (**b**) Isometric side-view of two regions denoted by dotted lines (width of line ROIs 200 nm), showing resolution of ring-like cross-profile. (**c, f**) Isometric side-view montages of 4 contiguous sections (section widths: 200 nm) corresponding to denoted regions in (a). (**d, g**) Scatter plots of the localizations from (c,f) aligned relative to the centroid. (**e**) Histogram of z-coordinates from (d) relative to the centroid, with inter-modal distance, gaussian fits, and standard deviations of subpeaks shown. (**h**) Radial distribution of the localizations from (g) relative to the centroid, with gaussian fit and fit parameters shown. (**i**) Axial color-coded 3D images of AlexaFluor647-labelled F-actin in COS-7 cells. (**j**-**l**) Inverted contrast side-view of regions denoted by red box in (i) and corresponding axial histograms (k, l). (**m**) Montage of axial color-coded images of actin networks from box region in (i), with the specified z-range indicated. Scale bars: 1 µm (a, m), 2 µm (i).

Next we imaged the actin cortex in COS7 cells, as labelled by phalloidin-conjugated to AlexaFluor 647 (**Fig. 3i**). Transverse axial sections across the cell (red box in **Fig. 3i, Fig. 3j**) reveals that ventral and dorsal actin cortex are clearly resolved. Axial histograms for two selected regions (arrowheads in **Fig. 3i**) are shown in **Fig. 3k** and **Fig. 3l**, with Gaussian fits showing standard deviation ranging from 9.7 nm to 21.91 nm, indicating that SiLM is capable of 10-nm axial precision even for dense filamentous meshworks. A montage of axial slices with 20-nm z-depth for the lamellipodial protrusion of the cell (white box in **Fig. 3i**) is shown in **Fig. 3m**. Distinctive actin profiles are observed in each axial section. Actin is observed to be largely excluded below 40 nm, with significant density beginning at z = 60 nm, consistent with previous measurements by iPALM^29^.

### Extended-Depth Interferometric Nanoscopy

Axial interferometry is conventionally limited in terms of imaging depth by phase-wrapping artefacts with a periodicity of λ/2n. To extend the imaging depth, previous iSMLM methods made use of non-periodic axial nanoscopic methods such astigmatic imaging^43^ or spline fitting to experimental PSF^12^. In SiLM, the extension of z-depth can readily be achieved simply by increasing the number of incidence angles. As described above and shown in **Fig. S1e**, the period of surface-generated standing waves in SiLM lengthens upon the increase in incidence angle. For 3-angle acquisition, phase-wrapping artefact is still present, with a period of about 250 nm for 639 nm excitation, as indicated by the CRLB analysis (**Fig. S2i-l**). To address this, 4-angle acquisition can be performed by a simple change in galvo control sequence (**Fig. 4a-e**). This provides an additional interference fringe (purple, **Fig. 4c**) with a period of around 500 nm, thereby resolving the phase-wrapping ambiguity. The 4-angle mode of SiLM allows nearly double the z-depth, extended to nearly 500 nm (**Fig. S2k-l**), as demonstrated in **Fig. 4b-e**, where microtubule filaments in immunostained COS7 cell are imaged. **Fig. 4d** shows the side-view of box region in **Fig. 4b**, whereas the isometric cross-section view of consecutive sections from **Fig. 4d** are shown in **Fig. 4e**, with the ring-like cross-profile of microtubules clearly resolved, reflecting that high axial precision is still maintained. Ongoing works are being pursued to harness the flexibility in incidence angle control for further axial precision enhancement and further imaging depth extension.

**Figure 4.**
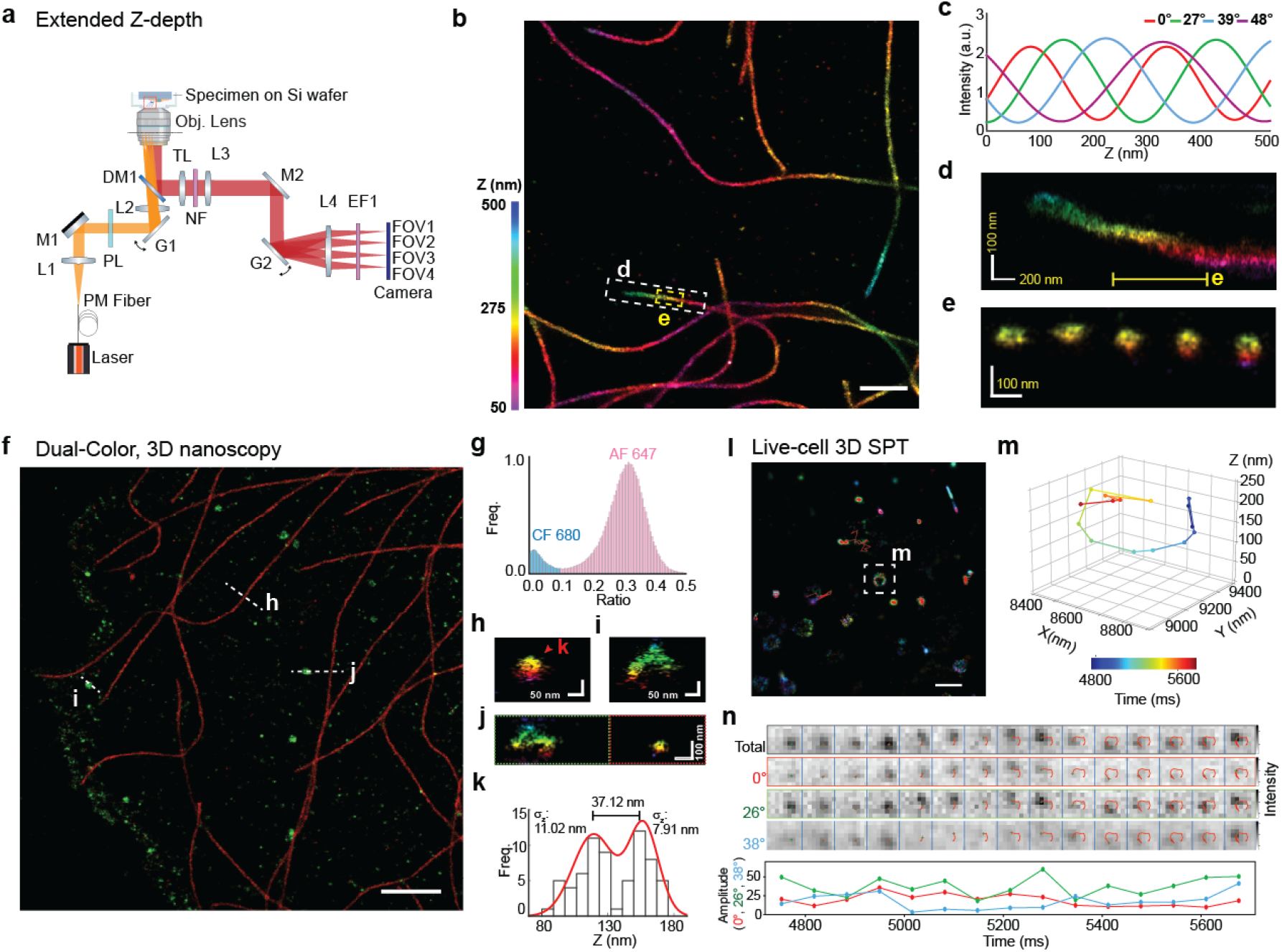
Versatility of SiLM. (**a**) Schematic diagram of 4-FOV acquisition for extended depth range axial interferometry. (**b**) Axial color-coded 3D image of AlexaFluor 647-immunolabelled microtubule filaments in COS-7 cells. (**c**) Simulated fluorophore intensity for θ = 0°, 27°, 39°, and 48° as a function of z-coordinate for SiO_2_ thickness of 481.4 nm and excitation wavelength of 639 nm. (**d**) Side-view profile of a microtubule from white box region in (b), showing an extended axial range. (**e**) Isometric side-view montage of 5 contiguous slices (section width: 200 nm) corresponding to yellow line in (d), showing ring-like cross-profiles of microtubules. (**f**) 2-color SiLM imaging of microtubule filaments (red, AlexaFluor 647 immunolabel) and clathrin-coated pits (green, CF680 immunolabel) in COS-7 cells. (**g**) Histogram of de-mixing ratio for AlexaFluor 647 (pink) and CF680 (blue). (**h-i**) Isometric side-view images from (f) with ring-like cross-profile of microtubule (h) from AlexaFluor 647 channel and hemispherical shape of clathrin-coated pit (i) from CF680 channel. (**j**) Isometric side-view along the indicated line in (f) depicting both a clathrin-coated pit (left) and a microtubule filament (right), with axial color-coding for CF680 and AlexaFluor647 channels, respectively. (**k**) Histogram of z-coordinates from (h) relative to the centroid, with inter-modal distance, gaussian fits, and standard deviations of subpeaks shown. (**l-n**) Live-cell 3D single particle tracking by SiLM, with trajectories of the membrane probe CAAX-SunTag_24_: tdStayGold-ScFv, overlaid on axial color-coded 3D image (l). (**m**) 3D temporal color-coded plot of a trajectory from boxed region in (l) depicting a circular motion around a vesicle. (**n**) Montages of raw images (488-nm excitation) at θ = 0°, 26°, and 38° and the summed raw images corresponding to trajectory in (l). Plot of intensity vs time for individual FOVs are shown on the lower panel. Scale bars: 1 µm (a, l); 2 µm (f).

### Dual-Color Interferometric Nanoscopy

For multi-color imaging, SiLM is particularly amenable to ratiometric demixing of emission spectra^48^. By an addition of a dichroic mirror and another camera (**Fig. S8a-b**), two fluorophores with similar excitation wavelengths but different emission spectra, such as AlexaFluor 647 and CF680, can be used simultaneously. Here, for each detected fluorophore, the ratio between the demixing photons (second camera) and the total photons can be used to distinguish between AlexaFluor 647 or CF680 (**Fig.4g, Fig. S8b**). 2-color SiLM imaging is shown in **Fig. 4f-k** with microtubules and clathrin-coated pits visualized by secondary antibodies conjugated with AlexaFluor 647 or CF680, respectively (see also **Fig. S9** for 2-color imaging of focal adhesions). The demixing ratio shown in **Fig. 4g** reveals two distinct peaks with the cut-off of 0.1 used to distinguish the fluorophore labels. Notably, SiLM maintains high axial precision in two-color mode, with the ring-like microtubule sections and the arc shape of clathrin-coated pits clearly resolved (**Fig.4h-k**).

### Live-cell 3D Single-molecule Tracking by SiLM

The complexity of iSMLM, especially 4Pi-based techniques^44,46,49^, entails a large number of mechanical degrees of freedom which amplify the influence of mechanical and thermal drifts. Live-cell imaging using iSMLM has thus far not been demonstrated. In contrast, SiLM involves a modular extension of a commercial fluorescence microscopy systems and thus can readily tap on their existing functionalities. As shown in **Fig. S7**, long-term focus stability can be readily maintained using commercial automatic focusing, and thus fiducial beads are unnecessary. This also enable live-cell SiLM imaging at physiological temperature for mammalian cells to be performed with active temperature and environmental control. As shown in **Fig. 4l**, the diffusion of the CAAX membrane probe conjugated with SunTag peptides and scFv -tdStayGold^50^ was monitored in live COS7 cells, using 488-nm excitation and the optimal incidence angles of 0°, 26 °, and 38°. **Fig.4m** shows the 3D trajectory of a molecule around a vesicle in box region in **Fig. 4l**, along with the montage of single-molecule intensity time-series for each FOV in **Fig. 4m-n**. Altogether such live-cell 3D single-particle tracking experiment further demonstrates the versatility of SiLM for biological imaging. Additional examples of SiLM with different experimental configurations can be found in **Fig. S8**.

### Accurate Mapping of Focal Adhesions Nanoscale Architecture by SiLM

The molecular specificity and nanoscale precision of super-resolution microscopy is ideally suited for mapping the nanoscale architecture of multi-protein complexes in cells^22^. In particular, the ultra-high axial precision of iSMLM techniques enable benchmark studies of compact structures such as integrin-based focal adhesions (FAs), which play central roles in cell migration and a broad range of mechanobiological processes^51^. However, while observations across multiple cell types have shown that proteins in FAs are axially stratified into multi-compartment architecture, there are also notable variability in the distribution of specific proteins between studies^29,52-55^. To what extent these reflect different experimental conditions or meaningful biological phenotypes has not been clear. Major challenges in systematically probing FAs at nanoscale level stem from the limited accessibility to iSMLM imaging capabilities. Toward addressing this with SiLM, we therefore sought to directly compare its performance in FA imaging against previous iPALM measurements^29^.

As shown in **Fig. 5**, we used human osteosarcoma cell line (U2OS) cultured on fibronectin-coated SiO_2_ wafer surface in comparable conditions to the earlier iPALM analysis^29^. However, instead of using overexpressed FA proteins fused to tdEos or mEos2 fluorescent proteins (FP), we probed for endogenous FA proteins by immunolabeling with AlexaFluor 647-conjugated secondary antibodies. We note that while FP fusion provides smaller linkage error, their ectopic overexpression can be perturbative. Hence, whether the localization of overexpressed FP fusion is representative of the endogenous localization, especially for multi-valent FA proteins whose individual fragments are capable of localizing to different FA compartments^29,56^, has major implication for biological interpretation. While the trade-off for immunolabeling is the larger linkage error, such labelling enabled us to probe the intrinsic distribution of endogenous proteins as well as biochemically active phosphorylated forms.

**Figure 5.**
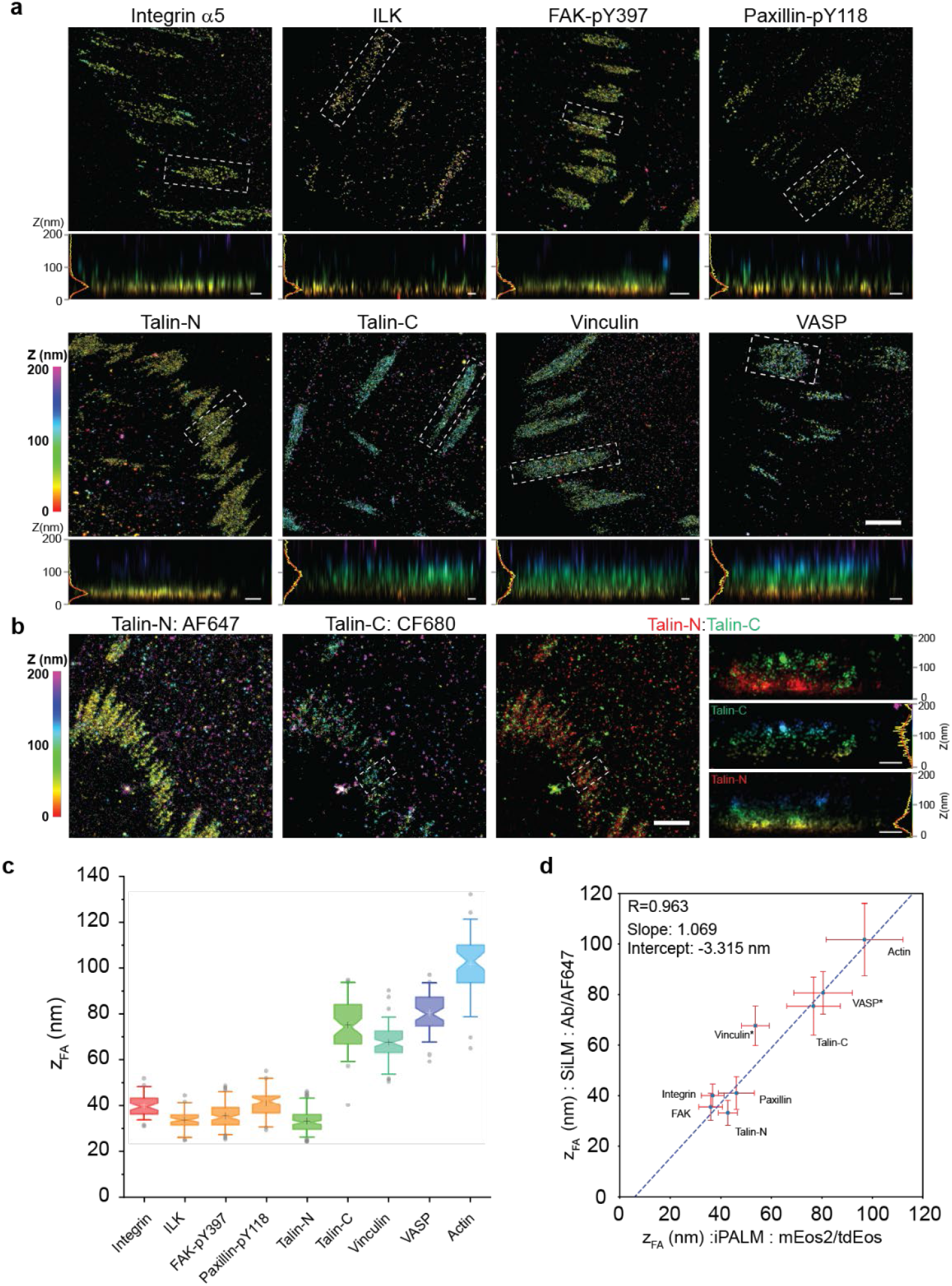
Accurate Mapping of Focal Adhesions Nanoscale Architecture by SiLM. (**a**) SiLM imaging of focal adhesion proteins in U2OS cells. 1^st^ and 3^rd^ rows: axial color-coded image of AlexaFluor 647-immunolabeled proteins. 2^nd^ and 4^th^ rows: side-view projection corresponding to boxed regions in upper panels, with histogram of z-position (yellow) and fit to single-gaussian (red) on the left. (**b**) 2-color SiLM imaging of talin1 orientation in focal adhesions. Axial color-coded image of talin1 immunolabeled against N-terminal epitope (left) or (2^nd^ from left) C-terminal epitope, with merged view shown in 2^nd^ from right panel (red: N-terminus, AlexaFluor 647; green: C-terminus, CF680). Side-view projection for boxed regions in left panels are shown on the rightmost columns with the merged red-green view on top. Sideview of C-terminal and N-terminal probes are shown in right-center and right-bottom panels, respectively, with z-position histogram (yellow) and gaussian fit (red) shown on the right. (**c**) Notched box and whisker plot depicting the distribution of median axial position of FA proteins (z_FA_), 5^th^ and 95^th^ percentiles indicated (see also Fig.S10a-b, Table S3). (**d**) Scatter plot comparing SiLM measurements using immunolabeling (y-axis) versus previous iPALM measurements using fluorescent protein fusions (x-axis)^29^. Average axial position of comparable proteins are plotted with the vertical and horizontal error bars corresponding to the standard deviation of SiLM and iPALM measurements, respectively. Linear regression and fit parameters are indicated. See also Figure S9-10. Scale bars: 2 µm (a, b, upper panels); 500 nm (a, b, sideview panels).

SiLM images of major FA proteins are shown in **Fig. 5a** with the side-view projection corresponding to highlighted FAs (boxed regions in upper panels of **Fig. 5a**) provided in the lower panels, along with their respective axial histograms and Gaussian fits (see **Table S3, Fig. S10a-b, and Fig. S10c-f** for actin). As in earlier analysis^29^, for each FA region, the axial distribution was fitted to a Gaussian to determine the peak position, z_FA_, and the width, σ_FA_, with the notched boxplots shown in **Fig. 5c** and **Fig. S10a-b**. We observed that integrin α5β1, integrin-linked kinase (ILK), the active Y397-phosphorylated form of Focal Adhesion Kinase (FAK-pY397), and the Y118-phosphorylated form of paxillin are localized in compact axial layers, with median z-positions of 39.48 nm, 33.70 nm, 34.78 nm, and 42.08 nm, respectively. These proteins were previously attributed to the membrane-proximal Integrin Signaling Layer (ISL)^29^. The localizations of the signalling active phosphorylated forms of FAK and paxillin corroborate their ISL designation while their axial localization are closely similar between SiLM and iPALM, as shown in **Fig. 5d.**

The ultra-high axial precision of iPALM has previously been used to resolve *in situ* molecular orientation of large proteins such as talin or vinculin^16,29,53,57^. Analogous experiment can be performed using SiLM for talin1, using antibodies against epitopes in the N- and C-terminal regions. This reveals the N-terminal domain of talin to be firmly anchored in the ISL zone (see also **Fig. S9a-c**) with median z-position of 32.64 nm, while the C-terminal domain of talin is elevated by 41.87 nm, positioned at median z-position of 74.51 nm. This indicates a highly polarized orientation with the C-terminal domain subjacent to the distribution of actin, as well as actin-associated protein such as VASP (**Fig. 5a, c**), and which is also in excellent agreement with prior iPALM measurements (**Fig. 5d**). We also performed 2-color SiLM imaging of N- and C-terminal epitopes of talin, respectively using AlexaFluor 647 and CF680 as fluorophores. As shown in **Fig. 5b**, this further corroborates the polarized orientation (**Fig. 5b, Fig. S9d-e**) with the C-terminus significantly elevated above the N-terminus, and establishing the ability of SiLM to resolve molecular orientation *in situ*.

Interestingly, for vinculin we observed a bimodal axial distribution in a significant proportion of FAs (**Fig. 5a, Fig. S10g-l**). Vinculin is an autoinhibitory mechanosensing protein, whose inactive form is localized to the lower ISL in FAs by interaction with phosphopaxillin, while activated vinculin is proposed to undergo upward translocation upon binding to talin and actin^53^. Indeed, when the axial distributions of vinculin are analyzed with bi-Gaussian fittings, distinct ISL and ARL populations are revealed (**Fig. S10a-b, g-k**), thus corroborating this model. We note that in previous iPALM experiments, a primarily unimodal distribution of vinculin was observed^29^. This is probably due to the use of overexpressed FP fusion proteins which may have differences in binding affinities compared to their endogenous counterparts. For example, prior to activation, vinculin is thought to be recruited to ISL via phosphopaxillin binding^53^. This may account for the unimodal and relatively lower axial position of Vinculin FP fusion as reported by prior iPALM measurements^29^, compared to the endogenous measurements using SiLM in this study (**Fig. 5d**). Notably, bimodal distribution of VASP, which appears to involve ISL and ARL subpopulations, are also observed by SiLM, (**Fig. S10a-b, l-p**). Indeed, VASP is a vinculin binding partner, whose nanoscale localization has been shown to depend on vinculin activation^15^. Whether these different nanoscale populations exhibit different biological activities is not currently known, but can be directly probed using antibodies specific to different phosphorylated forms.

Taken together, as shown in **Fig. 5d**, a correlation coefficient of R = 0.963 between SiLM measurements probing endogenous FA proteins and iPALM measurements using overexpressed FP fusions is obtained, indicating a very high degree of correspondence. This demonstrates that SiLM enabled a structural cell biology investigation of FAs with comparable performance to iSMLM techniques, but with the benefits of a significantly streamlined imaging and analysis process. Having established that endogenous FA proteins are organized into a stratified nanoscale architecture, SiLM can be readily deployed for further mechanistic investigation into how such nanostructural organization are organized and regulated in cells.

## Discussion

We demonstrated that leveraging surface-generated interference in a cost-effective, chip-based format enables highly accurate and precise 3D nanoscopic imaging through a simple extension compatible with most existing SMLM-capable microscopes. Our SiLM method offers several advantages. First, the axial (z) positions are internally referenced to the chip surface, providing absolute z-localization and eliminating the need for external fiducials or surface-adsorbed fluorophores as required in iPALM or related iSMLM methods. This minimizes sample-to-sample variation, reduces fiducial-induced artefacts, and dramatically improved robustness against mechanical drift^44^. Second, the single-objective configuration simplifies system construction, alignment, and calibration compared to 4Pi-based approaches^12,44,46,49^. Third, SiLM is modular and flexible, enabling diverse applications including multi-color, extended-depth, and live-cell 3D imaging. These advantages should benefit not only structural cell biology applications, such as mapping molecular architecture of supramolecular complexes in cells^22^, but also for scaling to automated high-throughput nanoscopy platforms.

Our demonstration of chip-based 3D interferometric nanoscopy by SiLM also opens up several promising directions for future development. Beyond basic Si/SiO_2_ substrates used here, alternative materials or multi-layer photonic structure may enable additional functionalities^30^. The current theoretical model, which assumes a homogeneous refractive indices in the sample, can be extended to address situations with heterogeneous refractive indices, thicker specimen, or when extremely high precision is desired. Speed and imaging area may be improved with faster switching devices, such as resonant scanner, and higher-performance camera^58^, including larger detectors. Additionally, recent studies have shown that linkage error, labelling density, and resonance energy transfer come into play for sub 10-nm precision^5,59^, but can be effectively addressed by labelling strategies such as RESI^6^. Since improvements in optical and bioconjugation approaches are largely orthogonal, SiLM can be seamlessly combined with these advanced labelling approaches for further resolution enhancement.

Our biological demonstration focused on integrin-based Focal Adhesions (FAs), whose benchmark 3D nanostructures have been probed primarily by iPALM^51^. We established a close correspondence between SiLM measurements of endogenous proteins and iPALM measurements of fluorescent protein fusions. This validates the accurate and high-precision 3D nanoscopic bioimaging capability of SiLM and corroborates the multi-layer architecture model of FAs^29^. Despite the central roles of FAs in mechanobiology, 3D nanoscopic mapping remains limited to mature adhesion states and fewer than 20 proteins across a small number of cell types, reflecting the constrained accessibility of iSMLM technologies^51,60^. With the streamlined design and operation of our chip-based approach, powerful 3D nanoscopic capability should now be broadly accessible for investigating the structural organization of not only integrin-based adhesions but also numerous other supramolecular complexes in cells.

## Methods

### Theoretical Foundation

The theorical basis for the formation of excitation standing wave by surface-generated interference above a reflective surface has been described previously^37-39^. Briefly, silicon is highly reflective in the visible spectrum while silicon oxide is highly transparent. Hence, a condition analogous to Lloyd’s mirror arises above a silicon wafer when direct laser excitation interferes with its reflection. The resulting standing wave, with fringe direction along the axial (z) dimension, can thus be used as a structured illumination pattern for axial nanoscopy. The intensity of such excitation field, *I(θ)*, can be described as follows:

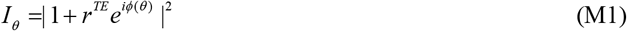

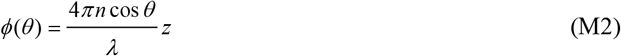

where *ϕ(θ)* represents the phase shift between the direct and reflected excitation, as experienced by a fluorophore positioned at a distance z from the surface of the silicon wafer substrate (defined as the reference z = 0 nm). In the context of the three-layer optical medium (silicon, silicon oxide, and specimen in aqueous buffer), the effective reflective coefficient *r*^*TE*^for s-polarized light can be expressed as follows:

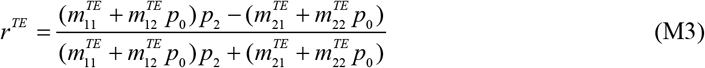

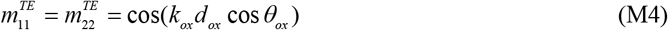

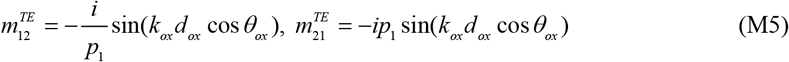

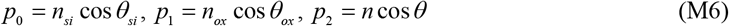

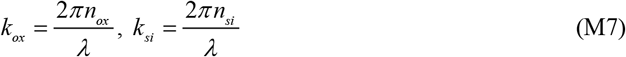

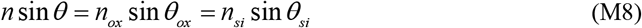

where *n, n*_*ox*_, *n*_*si*_represents the refractive indices in the sample, silicon oxide, and silicon layer, respectively. *d*_*ox*_is the thickness of the silicon oxide layer. With the definitions of transmission matrix *m*^*TE*^, and wavenumber k in Equations M4-M8, the effective reflective coefficient *r*^*TE*^can be calculated for each incidence angle, θ.

In SiLM experiments, the Si wafer is placed in aqueous imaging buffer in a glass-bottom dish. Therefore, it is important to account for the Fresnel transmission coefficient at the interface between the buffer and the glass. For *s*-polarized light, the transmission coefficient γ is given by:

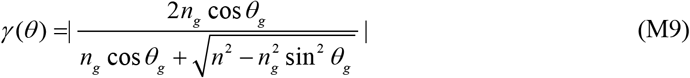

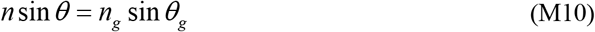

where *n*_*g*_and *θ*_*g*_are the refractive index and illumination angle at the glass-bottom dish, respectively. While the value of γ changes slowly for smaller angle, as θ_*g*_increases close to the critical angle, γ can increase substantially (**Fig. S1c**). Therefore, very large illumination angle is typically not chosen during actual experiments.

For a single-molecule fluorophore located at a distance *z* from the substrate surface, the fluorescence emission is expected to be proportional to *I*(θ)γ*(θ)*Hence, by modulating the excitation incidence angle *θ*, the fluorophore intensity is expected to scale proportionally with *I*(θ)γ*(θ)*. It can be shown that by acquiring a sufficient number (≥3) of intensity measurements at different values of *θ*, the parameter *z* can be solved, hence allowing axial localization analysis (see **Supplementary Notes I** for detailed theoretical analysis), while xy-coordinate determination can be performed similarly to conventional single-molecule localization microscopy (SMLM).

### Determination of optimal excitation angles

Theoretical analysis of imaging precision was performed using Cramer-Rao Lower Bound (CRLB) analysis as described in **Supplementary Notes I**. This enabled the calculation of imaging precision as a function of excitation wavelength, SiO_2_ thickness, and excitation incidence angles. A set of 3 incidence angles that enable the best axial localization precision values were optimized by global exhaustive multi-dimensional search to avoid local minima trap using GPU-accelerated computing code written in C and python. Tabulated values optimized for the axial range of 0-250 nm and a signal-to-noise ratio of 10 are shown in **Table S1**.

### Optical instrumentation

Our imaging system is based on a Nikon Eclipse Ti inverted microscope equipped with a motorized translation stage, a PerfectFocus system, and a live-cell imaging chamber (H301mini, OKO). For illumination, 405 nm, 488nm, 561nm, and 639 nm lasers (Coherent Cube Laser 1142279/AD 405nm/100 mW, Coherent Sapphire 488-200CW CDRH, Coherent Sapphire 561-200CW CDRH, CNI laser MRL-FN-639-400mW) were combined and directed through an AOTF (AOTFnC-VIS-TN1001/310905, AA-Optoelectronics), before subsequently coupled into a polarization-maintaining single-mode fiber (PM-S405-XP, Thorlabs). Detailed schematic of the setup is shown in **(Fig. 1a-b)** The fiber output is collimated and directed through the illumination galvo mirror G1 (GVS011, Thorlabs). A linear polarization plate (LPVISC 100-MP2, ThorLabs) is inserted after the galvo mirror to ensure that the excitation is *s*-polarized at the specimen. The lasers are then focused by a lens (AC 254-200A, Thorlabs) and directed by a dichroic mirror (Di03-R405/488/561/635-t1, Semrock Inc.) to the objective back focal plane from the back port of the microscope. The galvo plane is optically conjugated to the specimen plane. Objective lens used is either an oil-immersion Nikon CFI APO TIRF 60XC 1.49 NA or a Silicone oil-immersion Nikon CFI SR HP Plan Apo Lambda S 100XC SiI NA 1.35. Fluorescence emission from the specimen is collected by the objective lens in the standard epifluorescence configuration and routed to the left port of the Nikon microscope with the intermediate 1.5X magnification tube lens used.

For single-color imaging, at the output port a 4f system is constructed using 2 lenses (AC254-150-A, Thorlabs and AC508-200-A-ML, Thorlabs) with the detection galvo mirror G2 (Thorlabs GVS011) positioned at the intermediate Fourier plane. A notched filter (Notched Filter 405/488/561/635, Semrock Inc.) was used to reject stray excitation, and the FOV iris (SM1D12C, Thorlabs) is positioned at the immediate image plane after the left port of the microscope. A 2-inch-diameter lenses is employed as the second lens in the 4f system to reduce optical aberration in the point spread function during scanning. An EMCCD camera (iXon Ultra, Andor) is placed after an emission filter (ETF1: FF01-676/37, Semrock) to collect the fluorescence signal. The effective image pixel size is 133.33 nm. The control signals for AOTF, EMCCD camera triggering, and galvo mirrors are synchronized using National Instruments DAQ (Data Acquisition) devices (NI USB-6259 and NI PCI-6229) running a custom-written LabVIEW program. Schematic of the timing sequence is illustrated in **Fig. S3d**.

For dual-color imaging using the ratiometric approach, two fluorophores with a similar excitation wavelength but offset emission spectra (AlexaFluor 647 and CF680) are used since these allow single-wavelength excitation (**Fig. S8a-b**). An additional dichroic mirror DM2 (Multiband dichroic beam splitter: ZET 405/488/561/647rpc, Chroma) as well as an additional 4f system (AC254-150-A, Thorlabs, and AC254-200-A, Thorlabs) was added to detect fluorescence emission above 670 nm, using a second EMCCD camera (iXon Ultra, Andor). An emission filter EFT2 (FF01-661/20-25, Semrock) is positioned before the second camera. For the main optical path, the emission filter (EF1) is replaced with a multiband emission filter (ZET 405/488/561/647, Chroma). The ratio between second-camera photon count and the total photons were used to assign the labels to each localization coordinates.

For alignment, the excitation is first centred along the objective lens axis by adjusting the mirror (M1) in front of the illumination galvo G1 to align the output beam to a pre-calibrated spot on the room ceiling. To switch the illumination angle at the specimen plane, Analog voltage signals from the NI-DAQ device are applied to the illumination galvo G1. The angular movement of the illumination galvo (θθ_*galvo*_) is proportional to the applied voltage signal, and thus to achieve a specified incidence angle (θ) at the specimen in aqueous media, the following relationship is used:

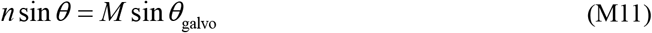

where M denotes the objective magnification, and *n is* the refractive index of specimen. After the optical system is built, additional angle calibration steps are performed as shown in **Fig. S3e-h**.

### Raw data acquisition

Before mounting each specimen, the excitation laser beam is manually centered by adjusting the mirror M1. The procedure for specimen mounting, focusing, and imaging site identification are similar to conventional fluorescence microscopy protocols. Once the desired imaging site is identified, the Perfect Focus System (PFS) was activated to continuously maintain focus. Subsequently, a calibration stack of 200 frames was collected in which the excitation incidence angle is kept constant for all field-of-view (FOVs). The calibration step was crucial for ensuring proper alignment and consistent image quality across each FOV. Following the calibration, acquisition was initiated whereby each FOV is synchronized with a specific excitation incidence angle; for example, FOV #1: 12°, FOV #2: 34 °, FOV #3: 48 ° for λ = 639 nm and ∼500 nm SiO_2_ thickness. For acquisition, the EMCCD camera is set to external trigger mode, with the frame time and exposure time set to 150 ms and 96 ms, respectively. The illumination galvo G1 cycles between three angles, with a dwell time of 0.9 ms and a switch time of 1.1 ms, while the detection galvo G2 is synchronized with the illumination galvo G1. The number of cycles, galvo dwell time, and galvo switch time can be adjusted in custom LabVIEW software. For AlexaFluor 647 and CF680 imaging, the cycles are repeated 16 times per each exposure duration. The timing diagram is shown in **Fig. S3d**. Between 10,000 and 50,000 raw frames are typically acquired (see **Table S2**). For live-cell tracking, the galvo dwell time is set to 8.9 ms, while the switch time is maintained at 1.1 ms. With the number of cycles set to one, the total exposure time is 30 ms. Accordingly, the frame time of the EMCCD camera is set to 66 ms.

### Calibration of the intensity between 3FOVs

As described above (**Fig. S1c**), the influence of transmission attenuation at the interface causes variations in the intensity of the illumination laser as a function of the incidence angle, which in turn leads to differences in the detected intensities. Additionally, as the detection galvo switches the image between different fields of view (FOVs) on the camera, each image traverses slightly different optical paths. Variations in these optical paths can introduce different optical aberrations, further contributing to intensity differences in single molecule images across the FOVs. To account for these intensity discrepancies, a calibration specimen is prepared using fluorescent beads deposited on a conventional glass coverslip. Subsequently, a stack of 200 images were acquired using the standard SiLM acquisition steps (**Fig. S4**), with synchronous control of the illumination and detection galvos across three illumination and detection angles. Following the data processing procedure outlined below, the number of photons collected in each FOV for each bead were extracted. The histograms of intensity ratios for the bead sample across the three channels were then analyzed. The median values of these histograms were used to determine the intensity ratios between the FOVs. These intensity ratios were then applied to accurately extract the axial positions.

### Data processing

To extract the 3D single-molecule coordinates, a procedure analogous to iPALM data analysis was used^10,44,49^. A detailed flowchart of the processing workflow can be found in **Fig. S4**. Briefly, following data acquisition, each FOV is cropped and transformed using the transformation matrix calculated from the calibration datasets. To determine the xy-coordinates, 2D localization analysis is performed as described above on the summation of all registered FOVs. Next, the intensity corresponding to individual FOV was determined by constrained least-square fitting, using the fitted xy-coordinate from the summed images as the fixed parameters. These yield a triplet of intensity values for 3-FOV acquisition, or a quartet for 4-FOV acquisition. Subsequently, the axial (z) coordinate was determined by fitting of the intensity sequences to the theoretical curves (Eq. M1-2; **Fig. 1; Supplementary Notes I**). For 2-color imaging, 2D localization analysis was performed for raw images acquired by the second camera. FOV registration was then performed relative to the 1^st^ camera dataset. Subsequently, the intensity corresponding to the same molecule in both cameras can be determined, and their ratio used to classify the fluorophore identity as described above.

To accelerate data processing, graphical processing unity (GPU) acceleration was implemented. Maximum likelihood and least-square fitting routines were implemented in CUDA and compiled in Microsoft Visual Studio 2017 to generate a dynamic link library (DLL) file. The CUDA code utilizes 128 threads per block and is called using the *ctypes* module in Python.

Drift correction was performed either using fluorescent fiducial beads deposited in the specimen when available or by the redundant cross-correlation (RCC) method described previously^61^. Reconstructed super-resolution images were generated by custom-written python code using the conventional 2D Gaussian representation with color indicating z-axis as indicated in individual figures^2,10^.

### Axial localization analysis

For the determination of z-coordinate from angle-dependent fluorescence intensity sequences, Least Squares Estimation (LSE) is used. Respectively, the objective functions can be expressed as:

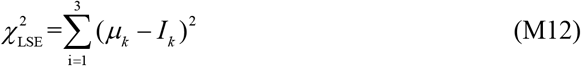

where µ_*k*_is the expected values of intensity as described by Eq. M1, and *I*_*k*_represents the single-molecule intensity observed at each incidence angle (i.e. each FOV). Two fit parameters are involved—the amplitude, *I*_*sig*_, and the axial position of the molecule, *z*. Levenberg-Marquardt (LM) method is used to solve the optimization problem. The descent direction for the parameter *x*_*j*_in LM method is given by the following equation:

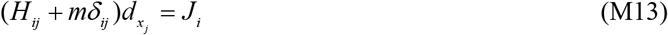

Here, the constant factor *m* and the diagonal matrix *δ*_*ij*_are used to ensure that the updates of the variables decrease the objective functions effectively. If the objective function does not descend, the value of *m* is increased (usually by a factor of 10) as needed. The matrices *H*_*ij*_and *J*_*i*_are defined by the following equations:

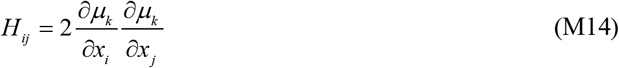

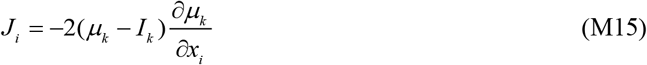

which only use the first derivative terms. In SiLM, the first derivative terms above can be defined as:

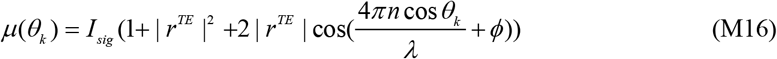

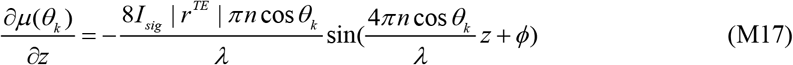

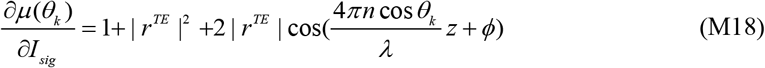

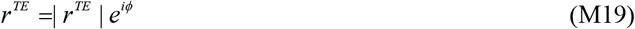

The complex term of the effective reflective coefficient *r*^*TE*^is represented by its amplitude and its phase angle are used as float variables. To overcome the local minima during optimization, several iteration routines can be calculated with variable starting axial positions, typically over a range of 0-300 nm at 50 nm interval. To expedite computation, GPU-accelerated code was written in C and compiled as dynamically-linked library (.dll) for interfacing with Python by *ctypes*.

### Silicon wafer functionalization

The imaging substrate used are 4-inch diameter silicon wafers (Bonda Technology) with nominal 500 nm thickness of thermal SiO_2_ layer. The precise oxide layer thickness of each wafer was determined using an ellipsometer (UV-VIS-VASE, JA-Woollam) at the Solar Energy Research Institute, National University of Singapore. A diamond-tipped pen was used to cut each wafer into 0.5 cm × 0.5 cm chips. Subsequently, the wafers were cleaned using acetone (Fisher Chemical A/0600/17, CAS: 67-64-1) and rinsed thrice in Mill-Q water. For chemical activation, the wafers were sonicated for 20 min in 1 M potassium hydroxide (KOH, Merck Millipore, catalog no. 22147, CAS: 1310-58-3), and then incubated for 1 h in 0.5% (3-aminopropyl) trimethoxysilane (Merck Millipore, catalog no. 281778, CAS: 919-30-2) in Milli-Q water. Wafers were then sonicated five times in water for 5 min each to remove excess (3-aminopropyl) trimethoxysilane, then incubated for 1 h in 0.5% glutaraldehyde in PBS, sonicated five times in water for 5 min and dried under nitrogen gas. Prior to use, wafers were UV-sterilized and incubated for 1 h at 37° with 10 µg/ml Fibronectin bovine plasma (Millipore, catalog no. f1141).

### Cell Culture

U2OS (Human osteosarcoma) and COS-7 (African green monkey kidney fibroblast-like) cells was obtained from American Type Culture Collection (Manassas, VA). Cells were adapted to and cultured in Glutamax DMEM supplemented with 10 % fetal bovine serum (FBS), 1 % sodium pyruvate, and 1 % penicillin/streptomycin solution (all from Life Technologies) in a constant humidity incubator set to 37°C and 5% CO_2_. On the day before imaging, cells were detached with 0.05 % Trypsin-EDTA solution (Life Technologies) and seeded at a density of ∼ 6,000 cells/cm^2^ on the fibronectin-coated thermal-oxide surface of Silicon Wafer described above. Cells were then allowed to grow on thermal oxide-containing side for 24–48 h in a 37 °C and 5% CO_2_ incubator before fixation.

### Immunolabeling

Cells were pre-fixed and permeabilized for 3 mins with 0.3% Glutaraldehyde (GA, Electron Microscopy Sciences, catalog no. 16200) and 0.2% Triton X-100 (Sigma-Aldrich, CAS: 9002-93-1) in PHEM (1x, PIPES 60 mM (Sigma-Aldrich, catalog no. p6757), HEPES 25mM (Sigma-Aldrich, catalog no. v900477), EGTA 10 mM (Sigma-Aldrich, catalog no. e3889), MgCl_2_ 2 mM (Sigma-Aldrich, catalog no. m1028), pH 7.4) buffer. Samples were then post-fixed for 15 mins using 4% Paraformaldehyde (PFA, Electron Microscopy Sciences, catalog no. 15710) and 0.2%Triton X-100 in PHEM, rinsed 3 times with PHEM. Fixative quenching was performed using 5 mg/ml sodium borohydride (Sigma-Aldrich, catalog no. 452882) in PHEM for 5 mins. After three washes with PHEM, cells were blocked by 3% BSA (Sigma-Aldrich, catalog no. a7906) and 0.1% Triton X-100 in PHEM for 30 mins, followed by staining with the primary antibodies overnight at 4^*°*^ in antibody dilution buffer (2% BSA and 0.02% Triton X-100 in PHEM). For actin staining, AlexaFluor 647 Phalloidin was used at 1:30 dilution. Cells were washed with washing buffer (1% BSA and 0.025% Triton X-100 in PHEM) 3 times for 5 mins per wash and stained with secondary antibodies for 60 mins at room temperature in antibody dilution buffer. Before imaging, samples were washed 3 times for 10 mins per wash with washing buffer.

Anti-α tubulin primary antibody was purchased from Abcam (1:200, catalog no. ab7291). Anti-clathrin heavy chain primary antibody was purchased from Abcam (1:200, catalog no. ab21679). Anti-Integrin-α5 primary antibody was purchased from BD Biosciences (1:200, catalog no. 555650). Anti-Phospho-Paxillin (Y118) primary antibody was purchased from Cell Signaling Technology (1:200, catalog no. 2541S). Anti-Phospho-FAK (Y397) primary antibody was purchased from Abcam (1:200, catalog no. ab81298). Anti-ILK primary antibody was purchased from BD Biosciences (1:200, catalog no. 611803). Anti-Talin-1(N-terminus, a.a. 139-433) primary antibody was purchased from Merck Millipore (1:200, catalog no. mab1676). Anti-Talin-1(C-terminus, a.a. 2491-2541) primary antibody was purchased from Abcam (1:200, catalog no. ab241096). Anti-Vinculin primary antibody was purchased from Sigma-Aldrich (1:200, catalog no. v9131). Anti-VASP primary antibody was purchased from Life Technologies (1:200, catalog no. ma5-14982).

AlexaFluor 647 Phalloidin was purchased from Life Technologies (1:200, catalog no. A22287). Donkey anti-mouse IgG (H+L) highly cross-adsorbed secondary antibody, Alexa Fluor 647 was purchased from Life Technologies (1:200, catalog no. A31571). Donkey anti-rabbit IgG (H+L) highly cross-adsorbed secondary antibody, Alexa Fluor 647 was purchased from Life Technologies (1:200, catalog no. A31573). Anti-Rabbit IgG (H+L), highly cross-adsorbed, CF660C antibody produced in goat was purchased from Merck Millipore (1:200, catalog no. sab4600453). Anti-Mouse IgG (H+L), F(ab)_2_ fragment, CF680 antibody produced in goat was purchased from Merck Millipore (1:200, catalog no. sab4600361). Anti-Rabbit IgG (H+L), F(ab)_2_ fragment, CF680 antibody produced in goat was purchased from Merck Millipore (1:200, catalog no. sab4600362). For calibration, dark-red fluorescent (660/680) FluoSpheres carboxylate-modified microspheres of 0.02 µm was purchased from Thermo Fisher Scientific(actual diameter: 0.027 µm, catalog no. F8783). The fluorescent microspheres were diluted 20,000 times, then incubated with the immunofluorescent stained samples for 15 minutes, and finally rinsed with imaging buffer. For drift characterization, functionalized gold nanorods were purchased from NanoPartz (C12-25-650-TC-PBS-50-1, catalog no. n2319). The functionalized gold nanorod was diluted 2,000 times, then incubated with the immunofluorescent stained samples for 15 minutes, and finally rinsed with the imaging buffer.

For imaging, the samples were first incubated with imaging buffer: 0.04g/ml glucose (1st BASE, catalog no. bio-1101-1kg), 50 mM Tris-HCl pH 8 (1st BASE, catalog no. buf-1416), 0.05 mg/ml glucose oxidase (Sigma, catalog no. g2133-250ku), 1 µg/ml catalase (Sigma, catalog no. c9322-5g), 75 mM MEA (Sigma, catalog no. 30070), and 7% glycerol (Fisher Chemical, catalog no. 10579570) in PBS (1x, Biowest, catalog no. L0615), and then inserted onto glass-bottom dish (35mm glass base dish, IWAKI, catalog no. 3970-035), such that the cell-containing side is facing downward. The imaging buffer was filled between the silicon wafer and the glass dish, with the excess imaging buffer absorbed by a paper towel. Specimens were then sealed with melted vaselin-lanolin-paraffin mixture described previously^62^.

### Live-cell imaging by SiLM

COS-7 cells were seeded at a density of ∼ 6,000 cells/cm^2^ on the fibronectin-coated thermal-oxide surface of Silicon Wafer. Cells were then allowed to grow on the thermal oxide-containing side for 24h in a 37 °C and 5% CO_2_ incubator. CAAX-SunTag plasmid (pHRdSV40-UP-24xGCN4-2xIFP2-CAAX-24xPP7, addgene, #134833) and scFv-staygold plasmid (the superfolderGFP coding sequence in the scFv-superfoldGFP (addgene, #60907) backbone was replaced with tdStayGold from the tdStayGold-pcDNA3 plasmid (RIKEN, RDB19609) to create a chimeric construct by gene synthesis (Epoch Life Sciences). Plasmids were transfected with Lipofectamine™ 3000 Transfection Reagent, then cells were incubated at 37 °C under 5% CO2 for 12 h. Cells were washed twice with DMEM (−) (no additives) and incubated with DMEM (−) (no additives) before imaging. The raw data were processed as described in the Data Processing section to obtain the three-dimensional coordinates of molecules in each frame. Subsequently, trajectories were established based on the distance between molecules in consecutive frames using the python library *trackpy* ^63^.

### Statistics and reproducibility

Plotting and statistical analysis for SiLM measurements (linear regression and uni-/bi-gaussian fitting) were performed using MATLAB (Naticks, MA), OriginPro software (Northampton, MA), or custom-written python codes. Detailed information on imaging parameters can be found in Supplementary Table S2. For FA protein axial position, data are presented as median, with tabulated analysis presented in Supplementary Table S3. All representative microscopy images are presented with quantification of the entire data set.

## Data and Code Availability

Data that supports the plots and figures within this paper and other findings of study are available from the corresponding author upon reasonable request. Processing codes are based on previously published solutions as described in Methods or Supplementary information. Codes for surface-generated interference calculation, CUDA codes for SiLM fitting and basic analysis pipeline are available on github repository: https://github.com/KanchanawongLab/SiLM. All other codes are available upon reasonable request.

## Acknowledgement

The authors thank the microscopy, information technology, nanofabrication, and open lab cores of the Mechanobiology Institute (MBI) for infrastructure support. The research is funded by Ministry of Education Singapore under Academic Research Fund Tier 2 (MOE-T2-1-124 and MOE-T2EP3-0124-0012 to P.K.), and Academic Research Fund Tier 3 (MOE-T3-2020-01 to P.K.), National Research Foundation Singapore under the Quantum Engineering Programme (QEP-P-7) and Mid-Sized Grant (NRF-MSG-2023-0001), and MBI intramural funding (P.K.). H.L. is supported by MBI Graduate Fellowship.

## References

1 Pawley, J. Handbook of biological confocal microscopy. Vol. 236 (Springer Science & Business Media, 2006).

2 Betzig, E. et al. Imaging intracellular fluorescent proteins at nanometer resolution. Science (New York, N.Y 313, 1642–1645 (2006).

3 Rust, M. J., Bates, M. & Zhuang, X. Sub-diffraction-limit imaging by stochastic optical reconstruction microscopy (STORM). Nat Methods 3, 793–795 (2006).

4 Hess, S. T., Girirajan, T. P. & Mason, M. D. Ultra-high resolution imaging by fluorescence photoactivation localization microscopy. Biophys J 91, 4258–4272 (2006).

5 Lelek, M. et al. Single-molecule localization microscopy. 1, 1–27 (2021).

6 Reinhardt, S. C. et al. Ångström-resolution fluorescence microscopy. Nature 617, 711–716 (2023).

7 Radmacher, N. et al. Molecular Level Super-Resolution Fluorescence Imaging. Annu Rev Biophys 54 (2025).

8 Pavani, S. R. et al. Three-dimensional, single-molecule fluorescence imaging beyond the diffraction limit by using a double-helix point spread function. Proceedings of the National Academy of Sciences of the United States of America 106, 2995–2999 (2009).

9 Huang, B., Wang, W., Bates, M. & Zhuang, X. Three-dimensional super-resolution imaging by stochastic optical reconstruction microscopy. Science (New York, N.Y 319, 810–813 (2008).

10 Shtengel, G. et al. Interferometric fluorescent super-resolution microscopy resolves 3D cellular ultrastructure. Proceedings of the National Academy of Sciences of the United States of America 106, 3125–3130 (2009).

11 Huang, F. et al. Ultra-high resolution 3D imaging of whole cells. 166, 1028–1040 (2016).

12 Bates, M. et al. Optimal precision and accuracy in 4Pi-STORM using dynamic spline PSF models. Nature methods 19, 603–612 (2022).

13 Backlund, M. P., Shechtman, Y. & Walsworth, R. L. J. P. r. l. Fundamental precision bounds for three-dimensional optical localization microscopy with Poisson statistics. 121, 023904 (2018).

14 Wang, W., Huang, Z., Wang, Y., Li, H. & Kanchanawong, P. Vortex Interference Enables optimal 3D Interferometric Nanoscopy. Physical review letters 134, 073802 (2025).

15 Bertocchi, C. et al. Nanoscale architecture of cadherin-based cell adhesions. Nature cell biology 19, 28–37, doi:10.1038/ncb3456 (2017).

16 Liu, J. et al. Talin determines the nanoscale architecture of focal adhesions. Proceedings of the National Academy of Sciences 112, E4864–E4873 (2015).

17 Balzarotti, F. et al. Nanometer resolution imaging and tracking of fluorescent molecules with minimal photon fluxes. Science (New York, N.Y, aak9913 (2016).

18 Gu, L. et al. Molecular resolution imaging by repetitive optical selective exposure. Nature methods 16, 1114–1118 (2019).

19 Cnossen, J. et al. Localization microscopy at doubled precision with patterned illumination. Nature methods 17, 59–63 (2020).

20 Gu, L. et al. Molecular-scale axial localization by repetitive optical selective exposure. 18, 369–373 (2021).

21 Jouchet, P. et al. Nanometric axial localization of single fluorescent molecules with modulated excitation. 15, 297–304 (2021).

22 Liu, S., Hoess, P. & Ries, J. Super-resolution microscopy for structural cell biology. Annu Rev Biophys 51, 301–326 (2022).

23 Dasgupta, A. et al. Direct supercritical angle localization microscopy for nanometer 3D superresolution. Nat Commun 12, 1180 (2021).

24 Bourg, N. et al. Direct optical nanoscopy with axially localized detection. 9, 587–593 (2015).

25 Szalai, A. M. et al. Three-dimensional total-internal reflection fluorescence nanoscopy with nanometric axial resolution by photometric localization of single molecules. Nat Commun 12, 517 (2021).

26 Thiele, J. C. et al. Isotropic three-dimensional dual-color super-resolution microscopy with metal-induced energy transfer. Science Advances 8, eabo2506 (2022).

27 Chizhik, A. I., Rother, J., Gregor, I., Janshoff, A. & Enderlein, J. Metal-induced energy transfer for live cell nanoscopy. Nature Photonics 8, 124–127 (2014).

28 Oleksiievets, N. et al. Three-dimensional multi-target super-resolution microscopy of cells using metal-induced energy transfer and DNA-PAINT. bioRxiv, 2024.2004. 2002.587536 (2024).

29 Kanchanawong, P. et al. Nanoscale architecture of integrin-based cell adhesions. Nature 468, 580–584 (2010).

30 Helle, Ø.I. et al. Structured illumination microscopy using a photonic chip. Nature photonics 14, 431–438 (2020).

31 Diekmann, R. et al. Chip-based wide field-of-view nanoscopy. Nature Photonics 11, 322–328 (2017).

32 Schnitzbauer, J., McGorty, R. & Huang, B. J. O. e. 4Pi fluorescence detection and 3D particle localization with a single objective. 21, 19701–19708 (2013).

33 Li, X. et al. Three-dimensional structured illumination microscopy with enhanced axial resolution. Nature biotechnology 41, 1307–1319 (2023).

34 Rodriguez, P. F. G. et al. MAxSIM: multi-angle-crossing structured illumination microscopy with height-controlled mirror for 3D topological mapping of live cells. Biophysical Journal 122, 450a (2023).

35 Mudry, E., Le Moal, E., Ferrand, P., Chaumet, P. C. & Sentenac, A. Isotropic diffraction-limited focusing using a single objective lens. Physical review letters 105, 203903 (2010).

36 Yang, X. et al. Mirror-enhanced super-resolution microscopy. Light: Science & Applications 5, e16134–e16134 (2016).

37 Braun, D. & Fromherz, P. J. P. r. l. Fluorescence interferometry of neuronal cell adhesion on microstructured silicon. 81, 5241 (1998).

38 Ajo-Franklin, C. M., Ganesan, P. V. & Boxer, S. G. Variable incidence angle fluorescence interference contrast microscopy for z-imaging single objects. Biophysical Journal 89, 2759–2769, doi:10.1529/biophysj.105.066738 (2005).

39 Paszek, M. J. et al. Scanning angle interference microscopy reveals cell dynamics at the nanoscale. Nature Methods 9, 825–827, doi:10.1038/nmeth.2077 (2012).

40 Paszek, M. J. et al. The cancer glycocalyx mechanically primes integrin-mediated growth and survival. Nature 511, 319–325, doi:10.1038/nature13535 (2014).

41 Park, S. et al. Immunoengineering can overcome the glycocalyx armour of cancer cells. Nat Mater 23, 429–438 (2024).

42 Carbone, C. B., Vale, R. D. & Stuurman, N. J. N. m. An acquisition and analysis pipeline for scanning angle interference microscopy. 13, 897–898 (2016).

43 Brown, T. A. et al. Superresolution fluorescence imaging of mitochondrial nucleoids reveals their spatial range, limits, and membrane interaction. Molecular and cellular biology 31, 4994–5010, doi:10.1128/MCB.05694-11 (2011).

44 Shtengel, G. et al. Imaging cellular ultrastructure by PALM, iPALM, and correlative iPALM-EM. Methods Cell Biol 123, 273–294, doi:10.1016/B978-0-12-420138-5.00015-X (2014).

45 Huang, F. et al. Ultra-High Resolution 3D Imaging of Whole Cells. Cell 166, 1028–1040, doi:10.1016/j.cell.2016.06.016 (2016).

46 Wang, J. et al. Implementation of a 4Pi-SMS super-resolution microscope. 16, 677–727 (2021).

47 Jouchet, P., Poüs, C., Fort, E. & Lévêque-Fort, S. Time-modulated excitation for enhanced single-molecule localization microscopy. Philosophical Transactions of the Royal Society A 380, 20200299 (2022).

48 Friedl, K. et al. Assessing crosstalk in simultaneous multicolor single-molecule localization microscopy. Cell Reports Methods 3 (2023).

49 Wang, Y. & Kanchanawong, P. Three-dimensional Super Resolution Microscopy of F-actin Filaments by Interferometric PhotoActivated Localization Microscopy (iPALM). J Vis Exp, doi:10.3791/54774 (2016).

50 Hirano, M. et al. A highly photostable and bright green fluorescent protein. Nat Biotechnol 40, 1132–1142, doi:10.1038/s41587-022-01278-2 (2022).

51 Kanchanawong, P. & Calderwood, D. A. Organization, dynamics and mechanoregulation of integrin-mediated cell-ECM adhesions. Nature reviews 24, 142–161, doi:10.1038/s41580-022-00531-5 (2023).

52 Stubb, A. et al. Superresolution architecture of cornerstone focal adhesions in human pluripotent stem cells. 10, 1–15 (2019).

53 Case, L. B. et al. Molecular mechanism of vinculin activation and nanoscale spatial organization in focal adhesions. Nature cell biology, doi:10.1038/ncb3180 (2015).

54 Xia, S., Yim, E. K. F. & Kanchanawong, P. Molecular Organization of Integrin-Based Adhesion Complexes in Mouse Embryonic Stem Cells. ACS Biomater. Sci. Eng 5, 3828–3842, doi:10.1021/acsbiomaterials.8b01124 (2019).

55 Kumari, R. et al. Focal adhesions contain three specialized actin nanoscale layers. Nat Commun 15, 2547 (2024).

56 Franco, S. J., Senetar, M. A., Simonson, W. T., Huttenlocher, A. & McCann, R. O. The conserved C-terminal I/LWEQ module targets Talin1 to focal adhesions. Cell motility and the cytoskeleton 63, 563–581 (2006).

57 Barnett, S. F. & Kanchanawong, P. Visualizing the ‘backbone’of focal adhesions. Emerging Topics in Life Sciences 2, 677–680 (2018).

58 Huang, Z. & Kanchanawong, P. Ultra high-speed single-molecule fluorescence imaging. Journal of Cell Biology 222 (2023).

59 Helmerich, D. A. et al. Photoswitching fingerprint analysis bypasses the 10-nm resolution barrier. Nature Methods 19, 986–994 (2022).

60 Chastney, M. R., Kaivola, J., Leppänen, V.-M. & Ivaska, J. The role and regulation of integrins in cell migration and invasion. Nature Reviews Molecular Cell Biology, 1–21 (2024).

61 Wang, Y. et al. Localization events-based sample drift correction for localization microscopy with redundant cross-correlation algorithm. Opt Express 22, 15982–15991, doi:10.1364/OE.22.015982 (2014).

62 Shin, W. D. et al. in Live Cell Imaging: A Laboratory Manual (eds Robert D. Goldman, Jason R. Swedlow, & David L. Spector) (Cold Spring Harbor Laboratory Press, 2010).

63 Crocker, J. C. & Grier, D. G. Methods of digital video microscopy for colloidal studies. Journal of colloid and interface science 179, 298–310 (1996).

